# Ischemia/Reperfusion Injury and Oxidative Stress Impair Cardiac Desmin Proteostasis

**DOI:** 10.1101/2023.05.09.540017

**Authors:** Zixiao Li, Seungho Jun, Krishna K. Singh, Patrick J. Calhoun, Gizem Keceli, Krishna Patel, Hikmet Kadioglu, Nazareno Paolocci, Giulio Agnetti

## Abstract

**Background:** Though acute mortality by myocardial infarction (MI) has declined in past decades, MI still represents one of the leading causes of heart failure (HF) development. We recently demonstrated the accumulation of toxic desmin aggregates in patients with HF of ischemic origin. Since desmin aggregates are toxic for the heart we aimed to test whether their formation can be induced by oxidative stress as a proxy for reperfusion injury, as well as addressing the effects of therapeutic strategies aimed at reducing desmin aggregation with cardiac oxidative stress.

**Methods and Results:** We demonstrate here that oxidative stress is able to induce desmin aggregation, acutely, in a cell-specific and dose-dependent fashion. We also show that elevation of *O*-linked β-*N*-acetylglucosamine (*O*-GlcNAc) prior to or after oxidative stress reduces the formation of toxic desmin aggregates and its pro-aggregating desmin post-translational modifications (PTM). In addition, we show for the first time a role for the transmembrane protease serine 13 (TMPRSS13) with desmin cleavage in response to oxidative stress while desmin’s single cysteine plays a protective role from I/R injury, which is independent of gain or loss of desmin function.

**Conclusions:** The proliferation of desmin PTM-forms (i.e., proteoforms) and its aggregation hallmark acute and chronic cardiac stress and result in both loss of and gain of desmin function. We report here two novel mechanisms that could be targeted for therapy to preserve desmin homeostasis and cardiac function in the acute settings of oxidative stress and reperfusion injury.

## INTRODUCTION

Approximately 1 million Americans experience myocardial infarction (MI) every year (1). In addition to the high, acute mortality rate (5-14%), the loss of cardiac tissue resulting from ischemia-reperfusion (I/R) injury has significant and long-term health implications that include the development of heart failure (HF) in ý30-40% of patients (2). These data underscore the need to develop novel therapies that can target acute and long-term adverse effects in patients presenting with MI.

Desmin is the main protein component of the intermediate filaments (IF) cytoskeleton in muscle cells and classically mediates structural and functional integrity (3). We demonstrated the accumulation of desmin-positive amyloid aggregates (pre-amyloid oligomers, or PAOs, short fibrils, and large aggregates) (4) similar to those observed in the brain of Alzheimer’s and Parkinson’s patients, in cardiac tissue extracts from various experimental models of HF and human specimens from HF patients (5, 6). In addition, we demonstrated that abnormal desmin aggregation which is facilitated by specific post-translational modifications (PTM, i.e. phosphorylation and cleavage) is toxic for cardiac cells (6). Our published data also suggest different dynamics of desmin aggregation in human ischemic vs non-ischemic HF (6). Specifically, while desmin forms PAOs (∼50 kDa) in both ischemic and non-ischemic HF, the accumulation of short amyloid fibrils (∼200 kDa) is more prominent with the former (6).

In this study, we address whether desmin aggregation can result from acute, oxidative stress, such as that observed with reperfusion injury. In addition to more established PTM, it was recently reported that the *O*-linked conjugation of a residue of β-*N*-acetylglucosamine with Ser and Thr residues, termed *O*-GlcNAcylation, plays a role in preventing protein misfolding during translation (7). Pharmacological elevation of *O*-GlcNAc levels protects rodents from cardiac I/R injury (8, 9). Using primary cultures of cardiac myocytes, we tested here whether pharmacological elevation of *O*-GlcNAc levels prevents desmin aggregation. We also measured the contribution of phosphorylation and cleavage with oxidative stress *in cellulo* and with I/R injury *ex vivo* to link the accumulation of desmin cleavage to acute, ischemic injury.

While excessive desmin misfolding is detrimental because of the gain of toxic function of desmin aggregates, desmin aggregation also contributes to desmin loss of function. This effect is due to the ability of desmin aggregates to propagate misfolding in a prion-like fashion (10). In addition, cleavage is a direct cause of desmin loss of function and ultrastructural disarray (11). Desmin can be cleaved by caspase-6 in the heart (12), and calpain-1 (13) and AAA-protease (14) in the skeletal muscle. In addition, we report here the dynamic interaction between desmin and the transmembrane protease serine 13 (TMPRSS13), with oxidative stress and I/R injury. To our knowledge this is the first evidence that TMPRSS13 is expressed in the myocardium at the protein level. Lastly, we address the involvement of the direct oxidation of desmin in our models of oxidative stress and we address the protective role for the oxidation of desmin’s single cysteine with global I/R injury. We performed these studies using redox-dead-mimetic desmin mutants (Cys to Ala or Ser) and to our knowledge, this is the first example in the literature showing that a change in PTM stoichiometry is sufficient to significantly affect functional outomes.

In summary, we tested whether exposure to oxidative stress as seen in both chronic and acute HF, induces desmin aggregation and its prodromic PTM (e.g., cleavage). In addition, we linked the proliferation of misfolded and modified desmin to cardiac oxidative stress and established that one of the ways by which *O*-GlcNAcylation protects from cardiac I/R injury is through the preservation of desmin proteostasis. While we report for the first time a pathophysiological role for TMPRSS13 in the heart, we also show that desmin’s single cysteine plays a protective role against oxidative stress through a mechanism different than the “simple” prevention of loss or gain of desmin toxic function.

## METHODS

### Cell Cultures, Protein Extraction and Quantitation

Cultured neonatal rat ventricular myocytes (NRVM) and fibroblasts (NRVF) were used for the *in cellulo* experiments. Cells were isolated as previously described (6) and treated with H_2_O_2_ at increasing concentrations (10, 50, 100µM) for ∼16 hrs to induce desmin aggregation and cell toxicity. The *O-*GlcNAcase inhibitor Thiamet G (TMG) was utilized to elevate *O*-GlcNAc levels either 3 hrs prior to or 30 min after treatment with H_2_O_2_. At the end of treatment cell cultures were quickly rinsed in ice-cold PBS, harvested in 25 mM HEPES pH=7.4 in PBS, completed with protease (Mini, Roche) and phosphatase (Phos-stop, Sigma) inhibitors, and centrifuged for 3 min at 5,000 rcf at 4 °C. The resulting cell pellets were snap-frozen in dry ice in ethanol and stored at −80°C. Cell toxicity was measured in 96-well plates (∼5×10^4^ cells/well) using a lactate dehydrogenase (LDH) cytotoxicity kit (Sigma-Aldrich, Cat # 11644793001). Absorbance was measured at λ=490 nm. Cell viability was measured in 96-well plates using the Vybrant MTT kit (Thermo, Cat # V13154) as per the manufacturer’s instructions. Absorbance was measured at λ=540 nm.

### Protein Extraction from Frozen Heart Tissue

Frozen heart tissue specimens were dissected using a cold razor blade in dry ice and precisely weighed (∼25 mg per piece) and individually transferred in 1.5 ml vials filled with 5 volumes (V/W) of ice-cold 25 mM Hepes pH=7.4 in PBS completed with protease (Mini, Roche) and phosphatase (Phos-stop, Sigma) inhibitors as per manufacturer’s instructions. Two chilled metal beads (Grinding Ball Stainless Steel 2 mm cat# 224550010, Retsch) were then added to each heart tissue sample. Homogenization was performed for 2 min at 28 Hz using a mixer mill (MM400, Retsch) and ice-cooled racks. The resulting tissue homogenates were quickly spun down using a minifuge and transferred to a magnetic rack (Thermo) to hold the beads in place while the homogenates were aspirated and transferred to new clean vials. Beads were rinsed with 3 volumes (V/W) of the homogenization buffer, and the washes were re-united with the corresponding homogenates and centrifuged for 15 min (4°C, 18,000 rcf). The resulting supernatants (cytosol-enriched, soluble fraction) were separated from the pellets (myofilament-enriched, insoluble fraction) and both fractions were snap-frozen in dry ice in ethanol and stored at −80°C until further processing.

### MAL-PEG Labeling

Cell lysates from cardiomyocytes treated with 10 or 50 µM H_2_O_2_ and controls were labeled with MAL-PEG (5kDa methoxy polyethylene glycol maleimide, Sigma) following a slightly modified version of a previously published procedure (15). Briefly, protein pellets from cardiomyocytes were dissolved in RIPA buffer (Sigma) containing 0.1% Triton X-100 and protease inhibitors (Sigma). Approximately 75 µg of protein/sample were incubated with 1 mM MAL-PEG in RIPA buffer containing 0.1% Triton X-100 at room temperature, in the dark, and under rotation for 2 hrs. After labeling, the samples were subjected to methanol/chloroform/water extraction to remove excess MAL-PEG and precipitate labeled and unlabeled proteins. The resulting protein pellets were dissolved in 40 µl of 1X Laemmli SDS sample loading buffer (Bio-Rad) completed with 50 mM DTT, heated to 90°C for 10 min, and analyzed by SDS-PAGE/western blot. Technical controls (i.e., duplicate samples without MAL-PEG labeling) were also collected and analyzed for reference.

### Transgenic Expression of Desmin Redox-Dead Mutants via AAV

Adeno-associated viral (AAV) vectors were obtained from Vector Builder. An AAV.PHP.s serotype was chosen because of its newly reported, improved cardiac tropism (16). The viral construct contained a cardiac troponin T promoter upstream of the N-terminally Myc-tagged mouse desmin sequence [NM_010043.2] encoding for two different redox-dead desmin mutants (Cys 332 to Ala or Ser; or CA and CS, respectively) or the wild-type sequence. Of note, only one cysteine is contained in the whole desmin sequence. The desmin gene was followed by an IRES bi-cistronic element enabling independent translation of nuclearly-localized cerulean under an SV40 promoter (cTnT>Myc/{mDes[NM_010043.2]*(C332A/S):IRES: SV40 NLS/Cerulean: WPRE).

About 8-weeks-old mice were, anesthetized using 4% isoflurane for general anesthesia and 0.5% proparacaine hydrochloride ophthalmic solution for local anesthesia and retro-orbitally injected as described by Yardeni T et al (Yardeni et al., 2011). The virus was diluted in sterile saline to a final concentration of 2 x 10^12^ GC/ml before injection (150 µl/mouse, n=5 each). Following the injection, mice were subject to cardiac echography weekly to monitor for signs of adverse remodeling. After ∼8 weeks susceptibility of the hearts to I/R injury was measured through a Langendorff apparatus.

### Ischemia-reperfusion Injury in Isolated (Langendorff-Perfused) Mouse Hearts

Hearts were collected from ∼12-week-old WT mice after mice received 500 IU Heparin i.p. 15 min before surgery and were then euthanized via cervical dislocation. Hearts were quickly excised and transferred to ice-cold Krebs-Henseleit buffer (118mM NaCl, 25mM NaHCO3, 11mM D-glucose, 4.7 mM KCl, 1.2mM MgSO4, 1.2mM KH2PO4, and 2.00mM CaCl2) (17). The aorta was cannulated within 2 min from the heart explant and transferred to a Langendorff apparatus for retrograde perfusion with Krebs-Henseleit buffer warmed to 37.5±0.5°C and equilibrated with 95% O_2_ and 5% CO_2_ at a rate of 4.5 ml/min through a peristaltic pump. Coronary perfusion pressure was kept at 80±5 mmHg through a pressure chamber placed 100 cm above the heart. A water-filled latex balloon was inserted into the LV to measure cardiac functions including heart rate, LV-developed pressure, diastolic pressure, the rate of LV pressure development, and pressure decay via an analog-digital interface (MP160 Biopac System Inc., CA). The I/R group was subjected to a baseline of 20 min. After baseline perfusion, hearts were subjected to 30 min global ischemia via stopping retrograde perfusion, followed by 120 min of reperfusion. The hearts were then decannulated and sliced cross-sectionally to obtain a ∼2 mm ring in the middle of the long axis. The rings were placed in cryomolds filled with Optimal Cutting Temperature (OCT) compound (Fisher) and rapidly frozen for histology while the remainder of the heart tissue was snap-frozen in liquid nitrogen for biochemistry.

### Western Blot Analysis

Cell pellets and myofilament-enriched fractions from heart tissue were heat-denatured in 1X LDS buffer (Thermo) completed with 20 mM DTT for 7 min at 95°C. Protein concentrations in the resulting samples were quantitated by the EZQ assay (Thermo). Volumes containing 10 µg of protein were resolved using SDS-PAGE (NuPAGE 3-12% gels, Life Technologies or 10% acrylamide Tris-Glycine, Bio-Rad for desmin fibrillar aggregates) and transferred to a nitrocellulose membrane (Bio-Rad, Cat #L002050A) using a power blotter (Thermo, PIERCE Power Station, Model 22838) as per the manufacturer instructions at 25 V and ∼1.3 Amp for 10 min. The resulting membranes were then blocked in 5% non-fat dry milk (Carnation) diluted in Tris-buffered saline (TBS) completed with 0.1% Tween 20 (TBS-T) for ≥30 min. Membranes were subsequently probed with primary and secondary antibodies as listed in Table 1 and rinsed 3 times (10 min ea.) in 5% milk/TBS-T between antibody incubations. After two additional rinses with TBS-T and one in TBS (10 min ea.), a fluorescent signal was recorded with a near-IR Odyssey scanner (LI-COR). Membranes were then stripped (1.5% Glycine,1% SDS, and 1% Tween 20 (pH=2.2) for 5 min followed by staining with direct blue 71 (DB71, Sigma) as per the manufacturer protocol, to assess equal loading and transfer performance.

**Table 1.** Antibody Information.

| Antibodies | Brand | Cat # | Species | Type | Dilution |
| --- | --- | --- | --- | --- | --- |
| CTD110.6 Anti-O-GlcNAc | (Comer et al., 2001) | N/A | Mouse | primary | 1:1,500 |
| Mouse anti-desmin clone DE-U-10 | Sigma Aldrich | D1033 | Mouse | primary | 1:10,000 (For WB); 1:500 (For PLA) |
| Rabbit anti-TMPRSS13 | abcam | ab59862 | Rabbit | primary | 1:1,500 (For WB); 1:200 (For PLA) |
| Rabbit anti-vimentin | Sigma Aldrich | PLA0199 | Rabbit | primary | 1:5000 |
| In Situ PLA Probe Anti-Rabbit PLUS | Sigma Aldrich | DUO92002 | Donkey | secondary | 1:5 |
| In Situ PLA Probe Anti-Mouse MINUS | Sigma Aldrich | DUO92004 | Donkey | secondary | 1:5 |
| Goat anti-Rabbit IRDye 800CW | LI-COR | 925-32211 | Goat | secondary | 1:15,000 |
| Goat anti-Mouse IRDye 680 RD | LI-COR | 926-68020 | Goat | secondary | 1:15,000 |

### Filter Assay Analysis

Samples de-natured as per western blot analysis were further diluted with 1X LDS buffer to get a final concentration of 0.05 µg/µl. Briefly, with this method a custom-sized nitrocellulose filter (0.22 µm cut-off, BioRad cat. #162-0112) and a vacuum manifold (BioRad Dot-Blot apparatus, Cat # 84BR-31798) is utilized to trap large protein aggregates as previously described (6). After equilibrating the membrane in TBS containing 2% SDS (TBS-S), 200 µL of each sample was filtered through the membrane using the vacuum manifold. After three washes with TBS-S, the membrane was probed as per western blot analysis.

### Quantification of Desmin Phosphorylation by Phos-tag Gels

Phos-tag (Wako, Japan) SDS-PAGE is a convenient method to measure protein phosphorylation which we first employed to study desmin (5). Briefly, the Phos-tag pendant (5 µM) is co-polymerized in 8% acrylamide, tris-glycine gels doped with 10 µM Mn^2+^ ions to create a positively charged gel matrix. The interaction between the positively charged gel matrix and the negatively charged phosphorylated protein residues results in delayed migration which in turn enables the separation of differently phosphorylated proteoforms. Denatured cell lysates containing 20 µg of protein/sample were used for these experiments and a gel containing everything, but the Phos-tag pendant was run in parallel using the same samples for optimization and QC (negative technical control). After separation, the gels were rinsed once with 1X transfer buffer (Bio-Rad) prior to transfer and processing as per western blot analysis. In Phos-tag gels, phosphorylated forms of desmin migrate at higher apparent molecular weight as a function of the number of phosphate groups (phospho-sites) per molecule.

### Proximity Ligation Assay (PLA)

Coverslips plated with NRVM (-/+10 and 50 μM H_2_O_2_ for 48 hr) were fixed with 4% paraformaldehyde (PFA, Fisher Scientific, Cat#15710) in PBS for 20 min at RT and permeabilized with 0.1% Triton X-100 (Acros, Cat#21568-2500) in PBS completed with 0.1% Tween 20 (PBS-T) for 10 min. After three washes in PBS-T, the coverslips were blocked with Duolink Blocking Solution (Sigma, Cat#DUO82007) for 1 hr. Coverslips were subsequently probed with primary (Desmin/TMPRSS13) and secondary antibodies in Duolink Ab diluent (Sigma, Cat#DUE82008) according to Table 1. After washing with 1X wash buffer A (Sigma, Cat#DUO82046) and incubation with 1X ligation buffer (Sigma, Cat#DUO82027) for 30 min at 37°C, the coverslips were further rinsed with 1X wash buffer A two times 5 min and incubated with 1X amplification buffer (Sigma, Cat#DUO82011) for 1 hr and 40 min at 37°C. Coverslips were then washed with 1X wash buffer B (Sigma, Cat#DUO82048) 2x for 10 min, 0.01X wash buffer B for 1 min, mounted using Duolink PLA Mounting Medium with DAPI (Sigma, Cat#DUO82040), and imaged using a Leica SP8 microscope with a 63x oil-immersion objective.

### Data and Statistical Analyses

For western blot analysis, the total desmin signal (densitometrical analysis determined through Image J using the digitized .tif image files) was normalized to the average of the same signal for controls (=100%) whereas individual proteoforms (i.e., full length, fragments and “dimers”) were expressed as a function of their relative percentage of the total desmin signal for a given lane.

One-way analysis of variance (ANOVA) followed by Tukey’s post hoc analysis was used to analyze western blot and filter assay experiments, while two-way ANOVA followed by Tukey’s post hoc analysis was utilized to analyze phospho-desmin re-distribution with phos-tag gels. Statistical analyses were performed using Prism 9.01 (GraphPad Software, San Diego, CA). For all experiments, the number of independent biological replicates used is indicated by *“n”.* Results were considered statistically significant for *P*≤0.05.

## RESULTS

### An *in cellulo* Model to Study Susceptibility of Cardiac IF to Oxidative Stress

Oxidative stress is an established disease contributor to both acute and chronic cardiac disease. To address the specific involvement of cardiac desmin aggregation with oxidative stress, we first compared IF remodeling in neonatal rat ventricular myocytes (NRVM) expressing mainly desmin, to fibroblasts (NRVF) which express the highly homologous, mesenchymal, type III IF, vimentin. Intriguingly, both protein sequences contain only one cysteine located in close proximity in the 2B coil domain (C333 for desmin and C328 for vimentin) and the primary cultures used in these experiments were obtained from the same hearts.

After subjecting NRVM/F to different types of oxidative stress we determined that treatment with 10µM H_2_O_2_ for 16 hrs was sufficient to segregate the two cell types in terms of viability, resulting in a ∼27% increase in fibroblasts survival compared to cardiac myocytes (*P*=0.0025, n≥14) by MTT (**Figure 1b**). A similar difference in survival was observed with 50µM H_2_O_2_ (∼31%, *P*=0.0005, n≥14) while exposure to 100µM H_2_O_2_ resulted in ∼50-70% dead in both cell types. These results oriented us to the use of 10 and 50µM H_2_O_2_ as ideal concentrations to segregate the two cell types in terms of survival from oxidative stress and were therefore followed further. Indeed, cell toxicity was not significantly different in the two cell types when treated with no and 10µM H_2_O_2_ while treatment with 50µM H_2_O_2_ elicited a ∼32% increase in LDH release in NRVM compared to NRVF (*P*≤0.0001, **Figure 1c**).

**Figure 1.**
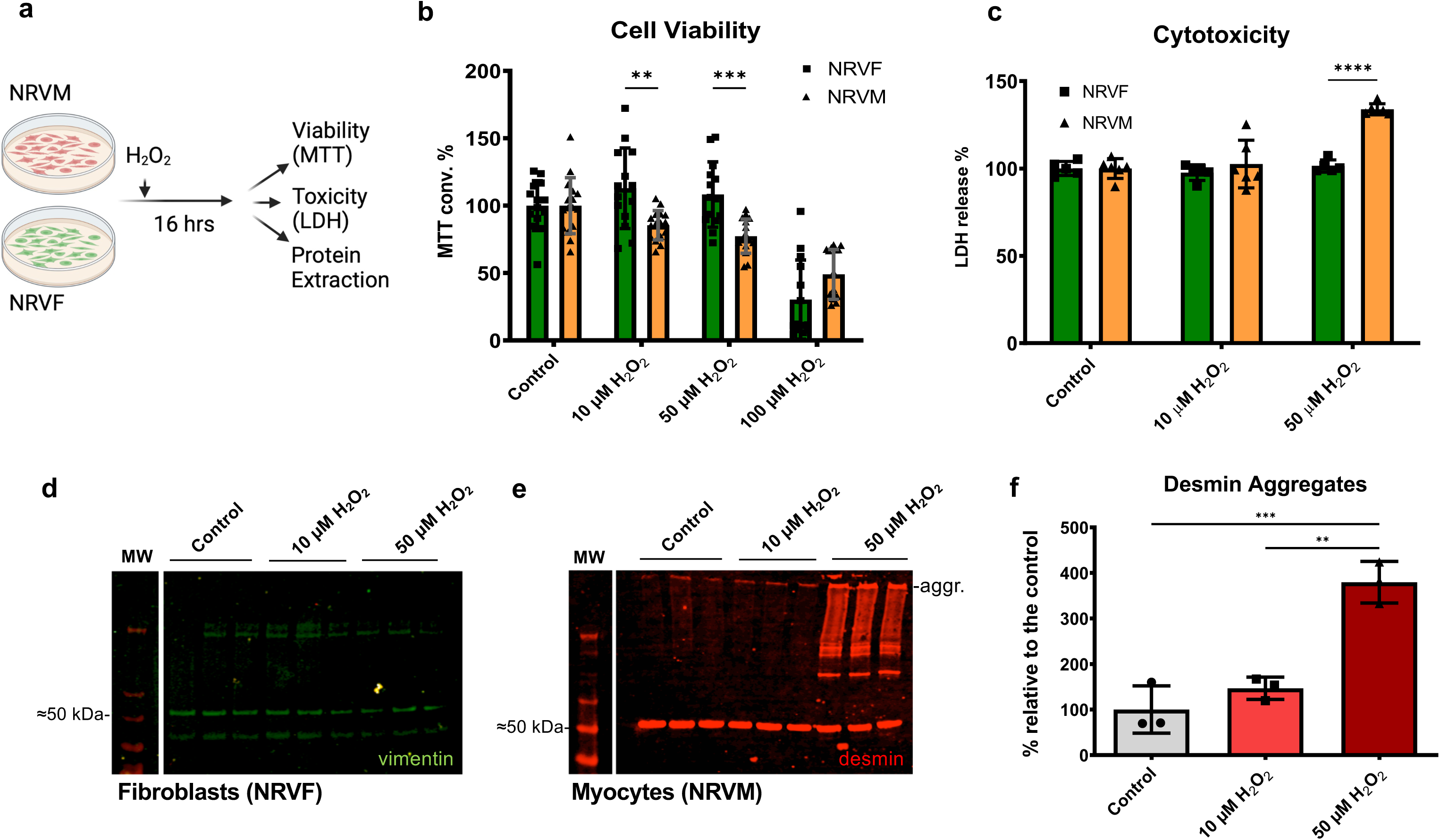
A Cellular Model to Address the Effects of Oxidative Stress on Cardiac IF Proteins. **a)** Experimental workflow. **b)** Normalized cell viability by MTT (*n*≥14) and **(c)** cell toxicity by LDH release (*n*≥5), in NRVM and NRVF treated with increasing concentrations of H_2_O_2_ for ∼16 hr. Representative western blot images of **(d)** vimentin in NRVF and **(e)** desmin in NRVM, and relative quantification of desmin aggregates at ∼200 kDa **(f**, *n*=3). Mean ± SD values are plotted. \*\*\*\**P*≤0.0001; \*\*\**P*≤0.001; \*\**P*≤0.01; by one-or two-way ANOVA followed by Sidak’s multiple comparison test. MW, Molecular weight markers; aggr., aggregates.

We recently proposed that IF protein aggregation could be used by cells to dissipate mechanical stress and that excessive aggregation could result in long-term toxicity (18). Increasing evidence points to a similar role for IF proteins in protecting cells from oxidative stress (19). As we previously reported a significant increase of short fibrillar desmin aggregates in human ischemic HF (6), we hypothesized that cardiac oxidative stress as seen with I/R injury (or chronic HF) could result in excessive desmin aggregation and that this effect could in turn be responsible for the increased risk of developing HF in ischemic patients. Therefore, we tested whether IF protein aggregation correlates with the susceptibility of cultured NRVM/F to oxidative stress by western blot, using Tris-Glycine gels (**Figure 1d-e**). While we could not detect any appreciable vimentin aggregation in fibroblasts exposed to increasing doses of H_2_O_2_ (**Figure 1d**), we could detect a ∼3-fold increase in desmin aggregation with 50 µM H_2_O_2_ at ∼200 kDa in NRVM (*P*<0.0001, **Figure 1e-f**). In all, these results highlight a cell-type, dose-dependent relationship between IF aggregation and cell toxicity/viability.

### Accumulation of Desmin Fragments with Oxidative Stress

One of the consistent findings in our published work is the accumulation of desmin cleaved forms with cardiac hypertrophy *in cellulo* and with heart failure, including humans, *in vivo* (5, 6, 20). We recently proposed that increased levels of desmin cleavage could reflect proteostatic insufficiency which ensues as a consequence of excessive desmin aggregation (18). Using an optimized approach to detect desmin cleaved forms through SDS-PAGE (see Methods section) we tested whether the accumulation of desmin cleaved forms is also induced by oxidative stress in NRVM. As shown above, treatment with 50 µM H_2_O_2_ induced marked cell injury in NRVM (**Figure 1c**). Cell toxicity in this cell type was accompanied by the marked accumulation of desmin cleaved products (∼3-fold, *P=*0.0023 compared to controls, **Figure 2a-e**) while the total and intact desmin levels were preserved (**Figure 2c-d**). Of note, a desmin proteoform at ∼100 kDa, compatible with a desmin dimer was also significantly increased with oxidative stress (∼8-fold, *P*<0.0001).

**Figure 2.**
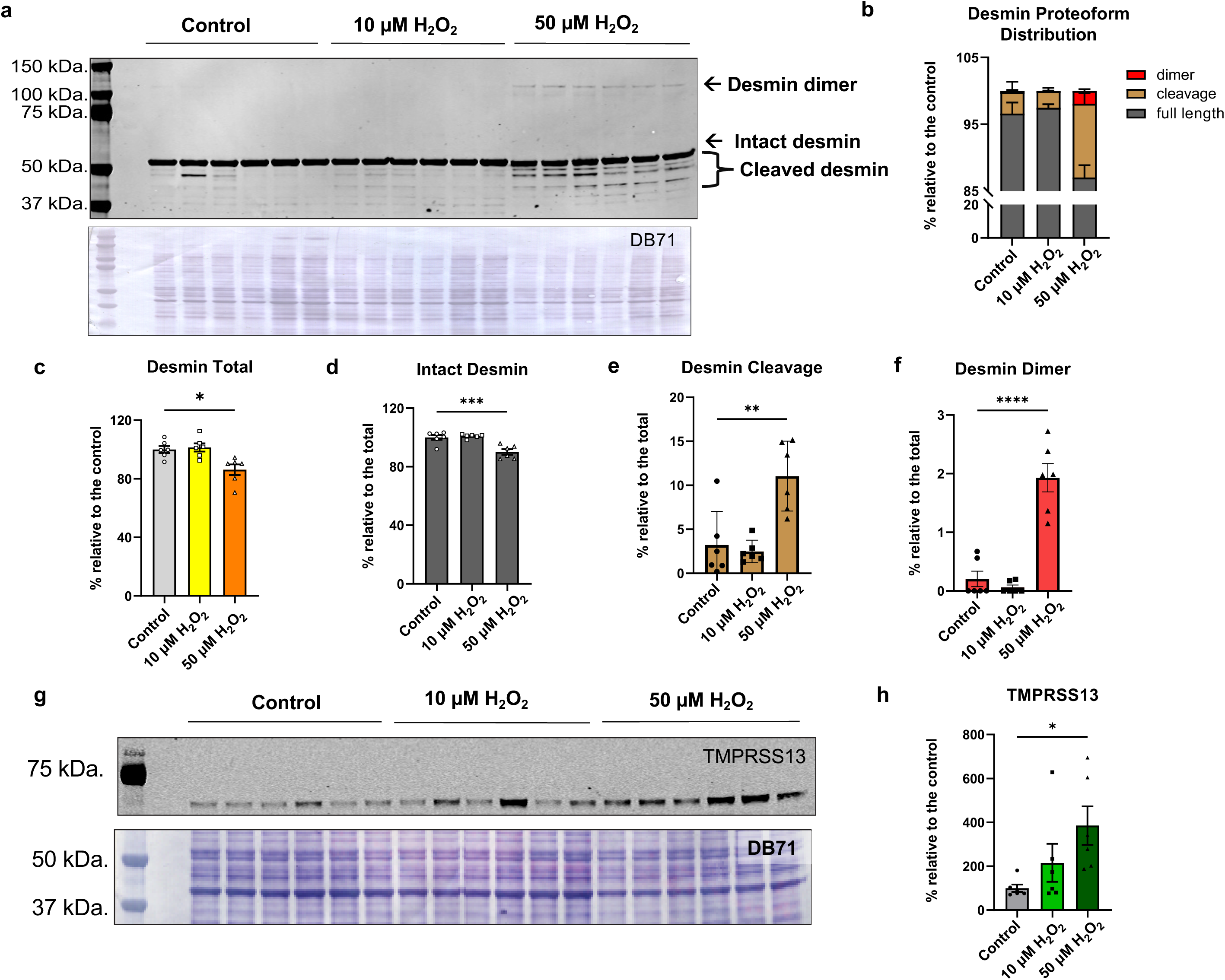
Desmin Cleavage Induce by Oxidative Stress Correlate with the Levels of TMPRSS13. **(a)** Representative western blot image for desmin (top) and direct blue 71 (DB71, bottom) for total protein staining. Quantitative measurement of desmin proteoform distribution **(b)**; total desmin relative to control **(c)**; full-length desmin relative to total desmin **(d)**; cleaved desmin relative to total desmin **(e)**; and dimerized desmin relative to total desmin **(f)**. Representative western blot image of full-length TMPRSS13 (top) and DB71 image for total protein staining (bottom) **(g)**; with relative quantitative measurement **(h)** (*n*=6). Mean ± SD is plotted. \*\*\*\**P*≤0.0001; \*\*\**P*≤0.001; \*\**P*≤0.01; \**P*≤0.05 by one-way ANOVA followed by Sidak’s multiple comparison test.

In a series of parallel experiments, we detected the interaction of the protease transmembrane protease serine 13 (TMPRSS13) in mice with HF vs. sham-operated controls by immuno-precipitation combined with mass spectrometry (MS, data not shown). Therefore, we measured the level of TMPRSS13 in NRVM and found a dose-dependent increase of TMPRSS13 levels with increased oxidative stress (**Figure 2g**). This increase in TMPRSS13 levels was significant with 50 µM H_2_O_2_ treatment (∼3-fold, *P*=0.0331, **Figure 2h**).

In short, these *in cellulo* data recapitulate the relationship between desmin misfolding and cleavage which we have previously reported *in vivo*, including in human patients with HF of both ischemic and non-ischemic origin (5, 6, 20), thus confirming the robustness and translational relevance of our *in cellulo* model. Based on sequence homology, TMPRSS13 is a membrane protease with the catalytic domain (C-term) exposed on the cell surface. For this reason, we sought to confirm that desmin and TMPRSS13 interact in our *in cellulo* model using a targeted approach.

### Oxidative Stress Increases Desmin-TMPRSS13 Interaction by PLA

Since both desmin fragments and TMPRSS13 levels were increased with oxidative stress in NRVM ± H_2_O_2_, we measured the direct interaction between desmin and TMPRSS13 using proximity ligation assay (PLA, Millipore Sigma). With this approach, the interaction between two proteins can be detected at the single molecule level in the form of puncti (21) and we thus confirmed increased desmin-TMPRSS13 interaction with oxidative stress, in a dose-dependent fashion. These data, linking desmin cleavage and TMPRSS13 levels and their direct interaction strongly suggest that TMPRSS13 contributes to desmin cleavage in our models. Very little is known of the function of TMPRSS13 in the heart and the biological significance of our newly reported interaction between TMPRSS13 and desmin with oxidative stress will be addressed in detail in follow-up studies.

### Differential Desmin Phosphorylation and *O*-GlcNAcylation Inversely Correlate with Susceptibility of Cardiac Myocytes to Oxidative Stress

We previously reported that the accumulation of mono-phosphorylated and cleaved desmin was associated with increased desmin aggregation in small and large animal models of HF as well as in tissue specimens from HF patients (5, 6). We also reported how mono-phosphorylated desmin at Ser-31 drives the formation of desmin PAOs which are toxic for cardiac cells (6). Therefore, we measured the levels of desmin phosphorylation using phos-tag gels (**Figure 4a**). Although treatment with 10µM H_2_O_2_ did not have a significant effect on the levels of mono-phosphorylated desmin, treatment with 50µM H_2_O_2_ significantly increased its levels compared to controls (by 1.07-fold, *P*<0.05, **Figure 4d**). Conversely, the levels of bi-, tri-, and tetra-phosphorylated desmin were more than 10-fold reduced by treatment with 50µM H_2_O_2_ compared to controls (*P*=0.0006, *P*=0.006, *P=*0.0002, respectively **Figure 4e-h**).

**Figure 3.**
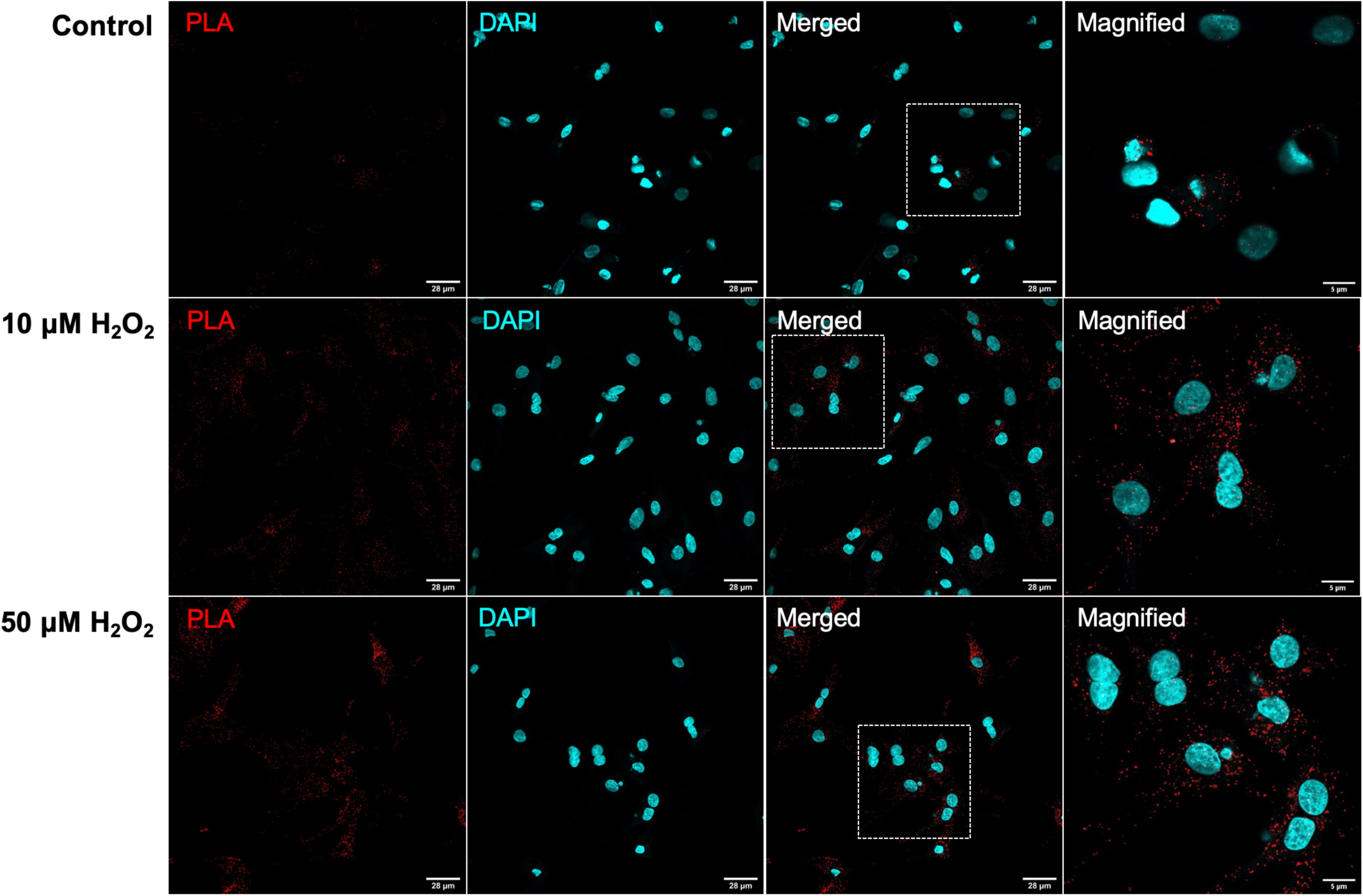
Proximity Ligation Assay for TMPRSS13 and Desmin. Interaction between TMPRSS13 and desmin in NRVM with increasing concentrations of H_2_O_2_. DAPI (in blue) was used to stain nuclei, whereas TMPRSS13-desmin interaction appears as red *puncti* (scale bars, 28 µm). Magnified images highlighting the interactions are also provided. (scale bars, 5 µm).

**Figure 4.**
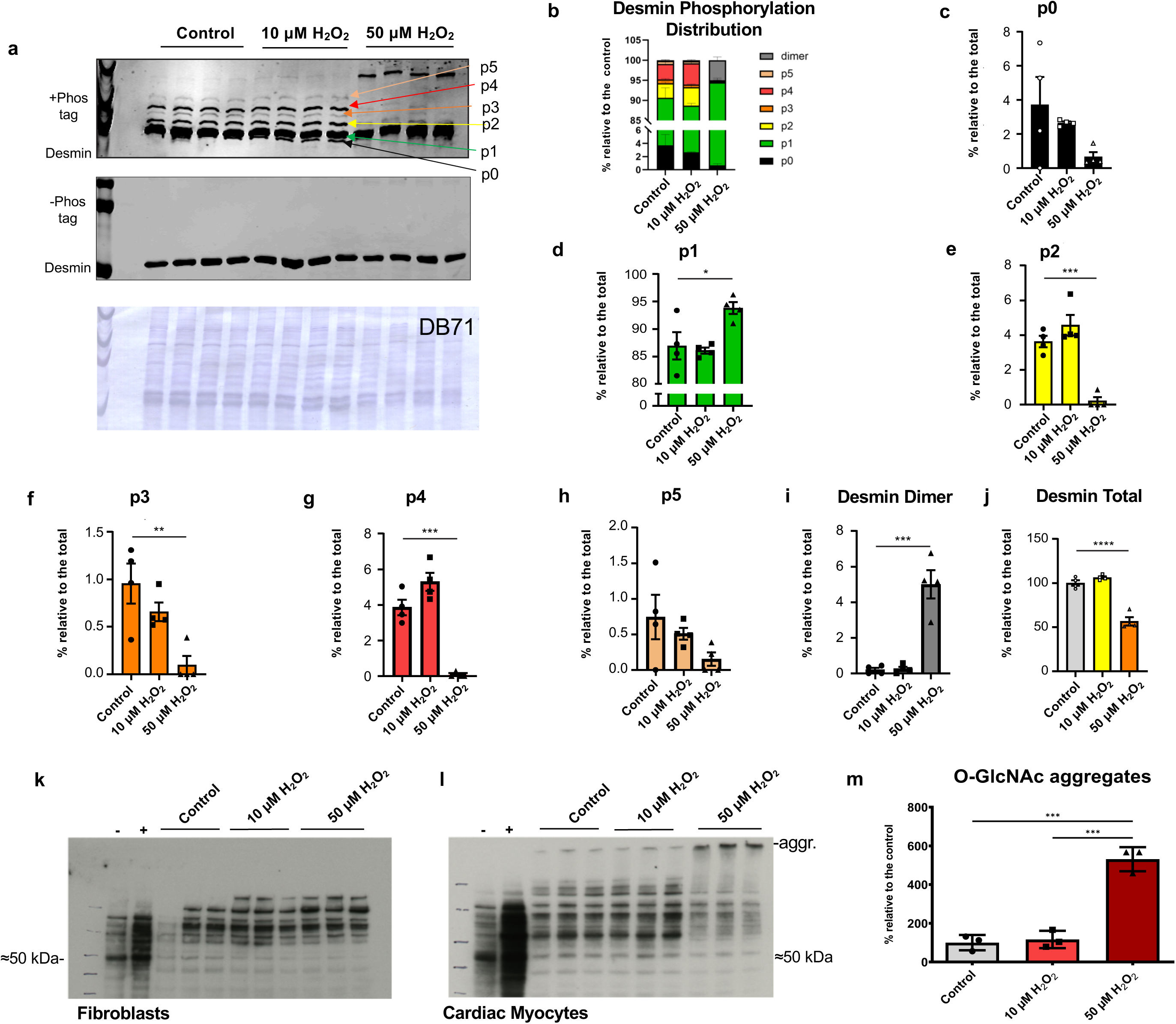
Effects of Oxidative Stress on Desmin Phosphorylation and O-GlcNAcylation. **(a)** Representative images of phos-tag (top panel) and phos-tag-negative control gels (mid panel) for desmin; direct blue 71 (DB71) image for total protein staining of phos-tag gel (bottom panel). Measurement of global **(b)** and individual phospho-desmin distribution **(c-h)** relative to total desmin. Quantitative measurements for dimerized desmin relative to total desmin **(i)**; and total desmin relative to control **(j),** are also provided (*n*=4). Representative western blot images of *O*-GlcNAc levels in fibroblasts **(k)**; and cardiac myocytes **(l). (m)** Quantitative measurement of O-GlcNAc-positive, desmin aggregates (*n*=3). Mean ± SD is plotted. \*\*\*\**P*≤0.0001; \*\*\**P*≤0.001; \*\**P*≤0.01; \**P*≤0.05 by one-way ANOVA followed by Sidak’s multiple comparison test.

These data are in agreement with our previously published data on large animal models and human specimens from HF patients (5) and support our proposed model where a biased accumulation of mono-phosphorylated desmin precipitates aggregation (22). Further, these data confirm that exposure of NRVM to H_2_O_2_ represents a convenient model to reproduce acutely the accumulation of desmin aggregates and its prodromic PTM that are observed in cardiac tissue specimens from HF patients.

Since the derivatization of Thr and Ser residue of intracellular proteins by *O*-GlcNAc (*O*-GlcNAylation) was recently shown to prevent protein misfolding during translation (7), we also examined whether *O*-GlcNAc would follow a similar molecular weight re-distribution to that of desmin aggregates. While no ∼200 kDa signal for *O*-GlcNAc was detected in neonatal fibroblasts used as negative controls (**Figure 4k**), a ∼5-fold increase in such signal, superimposed to that of desmin fibrillar aggregates at ∼200 kDa, was detected in NRVM (**Figure 4l**) treated with 50µM H_2_O_2_ (*P*<0.0001, **Figure 4m**, please compare to **Figure 1d**). An intriguing explanation for these findings is that cardiac myocytes attempt to prevent excessive desmin aggregation by “coating” fibrillar desmin with the large and hydrophilic *O*-GlcNAc moiety. Notably, we reported a similar accumulation of desmin fibrillar aggregates at ∼200 kDa with human ischemic HF (6). This combined evidence prompted us to test whether the established, protective effects of the pharmacological elevation of *O*-GlcNAc levels from I/R injury could be due at least in part to a reduction in desmin aggregation.

### *O*-GlcNAc Elevation Prevents the Cellular Cytotoxicity, Desmin Aggregation, and Desmin Cleavage Induced by Oxidative Stress

Pharmacological inhibitors of the *O*-GlcNAc-removing enzyme, *O*-GlcNAcase (OGA), protect against the detrimental effects of I/R injury (23, 24). However, the mechanisms underlying such protection are unclear. Here, we tested whether the protective effects of OGA inhibition via the small molecule Thiamet G (TMG) with oxidative stress are due, at least in part to the normalization of pathogenic desmin proteoforms (e.g., cleaved) and the reduction of its aggregation.

We first tested these hypotheses in NRVM treated with increasing doses (2 and 5 µM) of TMG 3 hrs prior to oxidative stress (50 µM H_2_O_2_). Inhibition of OGA through administration of TMG resulted in the dose-dependent increase of *O*-GlcNacylated proteins in NRVM (**Supplementary Figure 1**). In addition, treatment with TMG 3 hrs prior to oxidative stress was sufficient to reduce desmin cleavage in a dose-dependent fashion (**Figure 5a-c**). Specifically, accumulation of desmin fragments was reduced by ∼37% with 2 µM TMG (*P*<0.0001, n=3) and by ∼44% with 5 µM TMG (*P*<0.0001, n=3) when compared to treatment with 50 µM H_2_O_2_ alone (**Figure 5b**). Cytotoxicity, as measured by LDH release, was also markedly reduced (∼56%) in NRVM pre-treated with 5 µM TMG compared to 50 µM H_2_O_2_ alone (*P*=0.0117, n=3, **Figure 5d**), confirming a protecting role of *O*-GlcNAc elevation in our cellular model of oxidative stress. Lastly, pre-treatment with 5µM TMG reduced the levels of desmin aggregates by ∼two-fold by filter assay (*P*<0.0001, n=3, **Figure 5e-f**). This combined evidence supports the notion that the prevention of desmin cleavage and aggregation contributes to the protective effects of *O*-GlcNAc elevation in response to oxidative stress.

**Figure 5.**
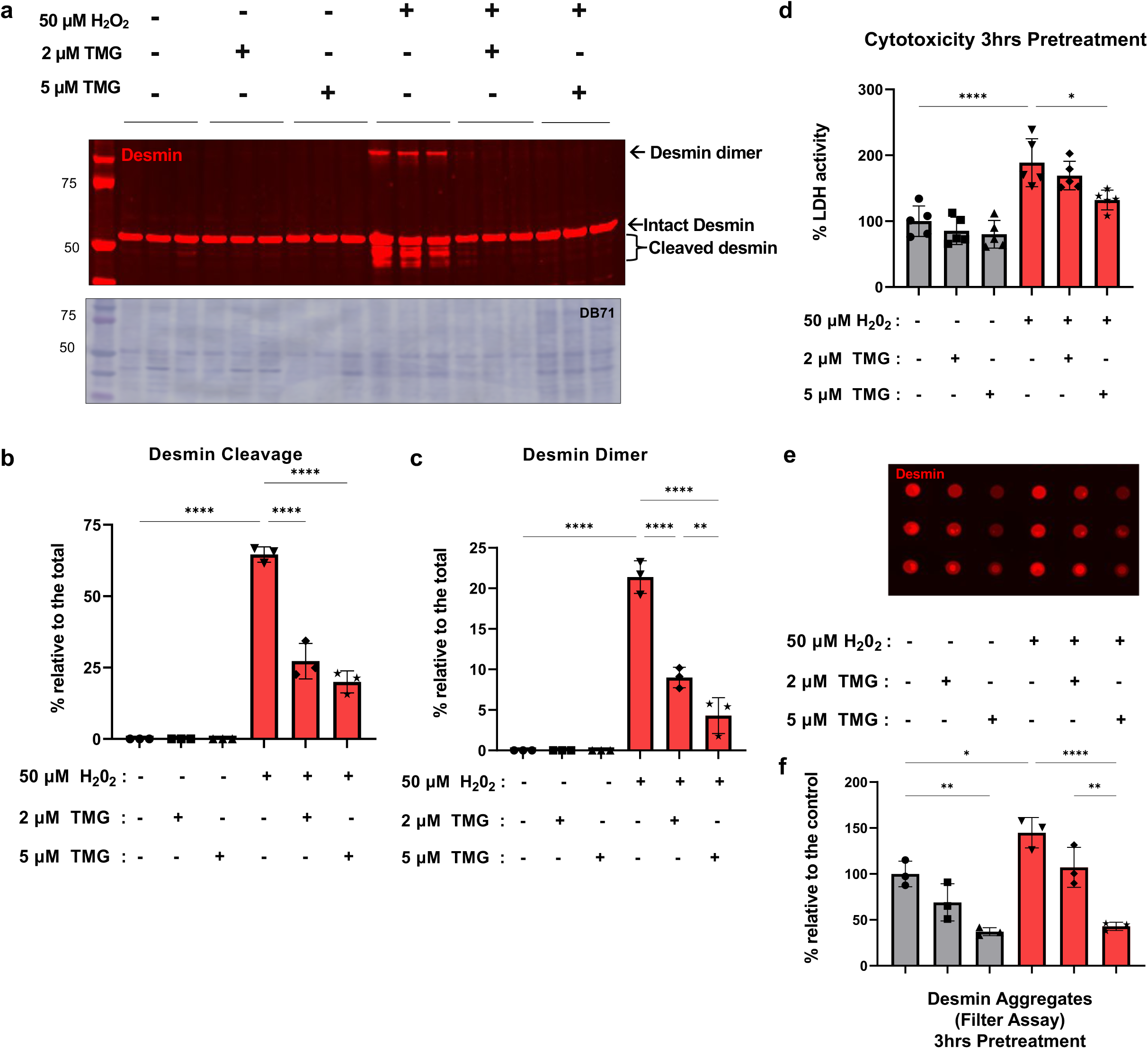
*O*-GlcNAc Elevation by TMG Prior to Oxidative Stress Preserves Desmin Homeostasis and Reduces Cell Toxicity. **(a)** Representative western blot image for desmin (top) and DB71 for total protein staining (bottom) of NRVM pre-treated with 2µM and 5µM TMG for 3 hrs before exposure to 50µM H_2_O_2_. Quantitative measurements of desmin cleavage relative to total desmin **(b)**; desmin dimers relative to total desmin **(c)**; and cytotoxicity by LDH release **(d)**. Representative image of a filter assay for desmin aggregates **(e)** and relative measurement of desmin aggregation ± 2 and 5 µM TMG/50µM H_2_O_2_ relative to controls **(f)** (*n*=3). Mean ± SD is plotted. \*\*\*\**P*≤0.0001; \*\**P*≤0.01; \**P*≤0.05 by one-way ANOVA followed by Sidak’s multiple comparison test.

To test the potential translational value of *O*-GlcNAc elevation with oxidative stress, we also administered TMG 30 min after oxidative stress with 50 µM H_2_O_2_ (**Figure 6**). Indeed, 30 min posttreatment with 5µM TMG was able to reduce desmin cleavage by ∼30% (*P*<0.0001, n=3, **Figure 6b**) compared to the controls. This trend was mirrored by a ∼50% reduction in LDH release (*P*=0.048, n=3, **Figure 6d**), and by more than 90% reduction in desmin aggregation (*P*=0.0002, n=3, **Figure 6e-f**).

**Figure 6.**
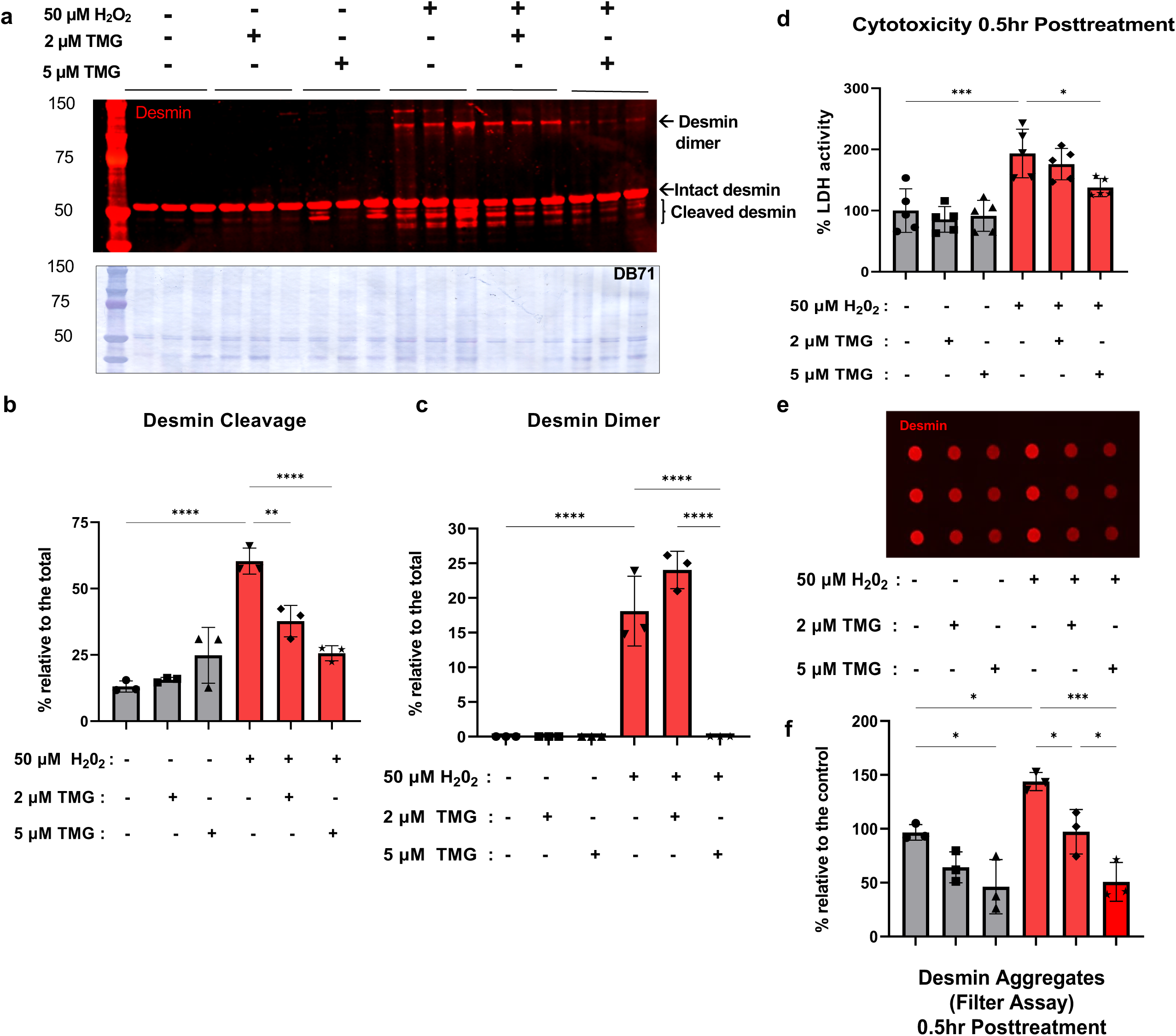
*O*-GlcNAc Elevation by TMG Following Oxidative Stress Preserves Desmin Homeostasis and Reduces Cell Toxicity. **(a)** Representative western blot image for desmin (top) and DB71 for total protein staining (bottom) of NRVM post-treated with 2 and 5 µM TMG 30 min after exposure to 50 µM H_2_O_2_. Quantitative measurements of desmin cleavage relative to total desmin **(b)**; desmin dimers relative to total desmin **(c)**; and cytotoxicity by LDH release **(d)**. Representative image of a filter assay for desmin aggregates **(e)** and relative measurement of desmin aggregation ± 2 and 5 µM TMG/50µM H_2_O_2_ relative to controls **(f)** (*n*=3). Mean ± SD is plotted. \*\*\*\**P*≤0.0001; \*\*\**P*≤0.001; \*\**P*≤0.01; \**P*≤0.05 by one-way ANOVA followed by Sidak’s multiple comparison test.

In all, this combined evidence suggests that the increase in desmin aggregation which results from the enrichment of cleaved desmin forms that we reported in chronic models of heart failure, can be corrected via pharmacological elevation of *O*-GlcNAc before or shortly after cardiac oxidative stress. Besides the direct implications to the mechanisms of *O*-GlcNAc-mediated cardioprotection, these data suggest that NRVM subjected to oxidative stress could represent a convenient platform to screen small molecules aimed at limiting the toxic effects of excessive desmin aggregation.

### Accumulation of Desmin Cleaved Forms is Induced by Ischemia/Reperfusion

To establish the relevance of desmin post-translational re-modeling and aggregation and ischemia-reperfusion (I/R) injury in the whole heart, we subjected 12-week-old C57BL/6 mice to I/R injury using the Langedorff method (15)(**Figure 7a**).

**Figure 7.**
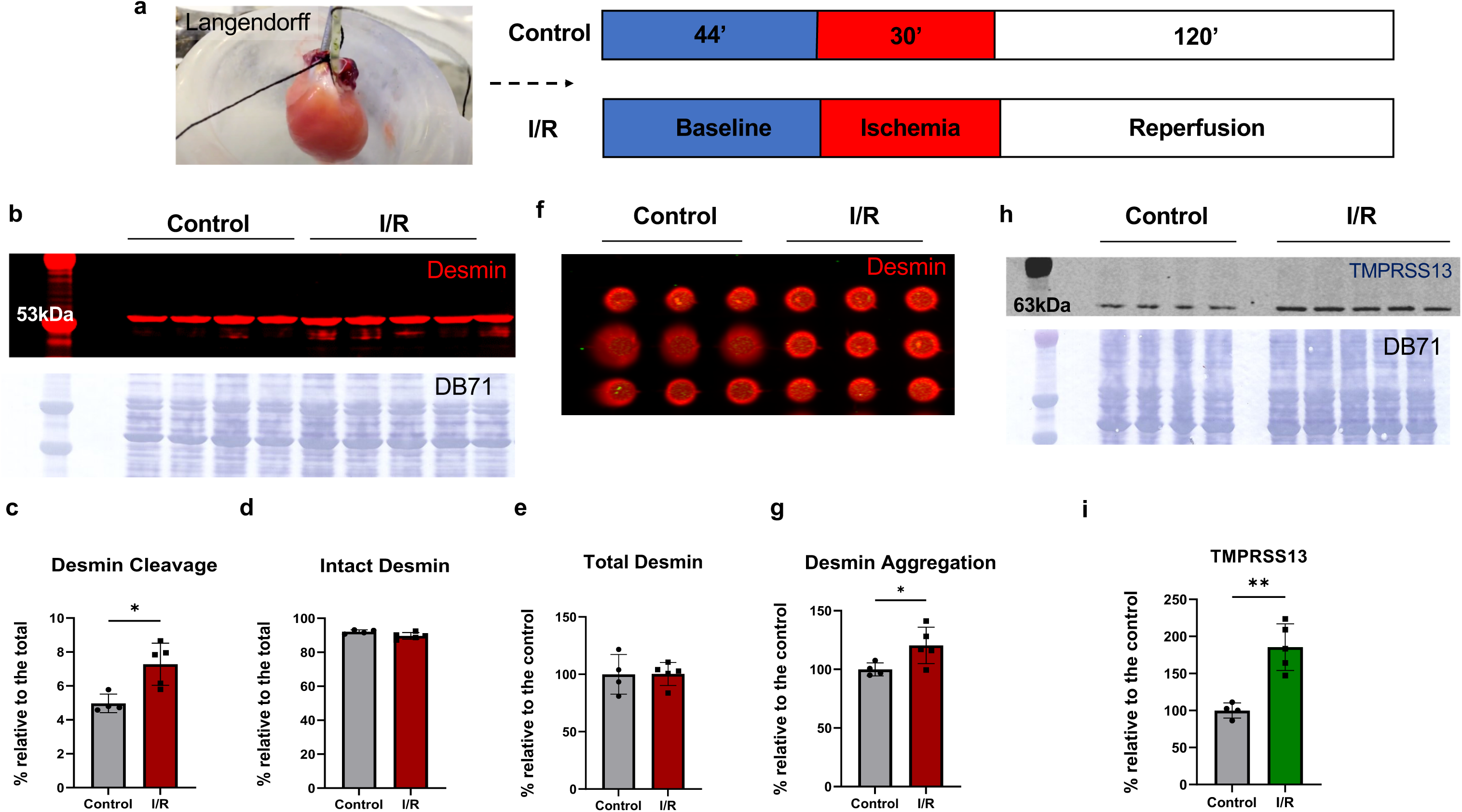
I/R Injury Increases Desmin Cleavage, Aggregation, and TMPRSS13 Levels. **(a)** Schematic of the I/R injury (top) and control (bottom) protocols used for hearts isolated according to Langendorff. **(b)** Representative western blot images for desmin with I/R and controls (top); and DB71 for total protein staining (bottom). Quantitative measurement of cleaved **(c)**; and intact desmin **(d)** relative to total desmin in each lane; and total desmin relative to control **(e)**. Representative filter assay image of desmin aggregation **(f)**; and relative measurement of desmin aggregation with I/R injury vs. controls **(g)**. Representative western blot image for full-length TMPRSS13 with I/R and controls (top); and DB71 for total protein staining (bottom) **(h)**; and relative measurement of full-length TMPRSS13 with I/R injury vs. controls **(i)** (*n*≥4). Mean ± SD is plotted. Mean ± SD is plotted. \*\**P*≤0.01; \**P*≤0.05 by student’s t-test.

Similar to NRVM subjected to treatment with H_2_O_2_, desmin cleavage was ∼46% elevated in I/R when compared to control mice (*P*=0.0109, **Figure 7b-c**). Of note, while there was no difference in the levels of total and intact desmin by western blot analysis (**Figure 7d-e**), desmin aggregation was ∼20%-increased in I/R compared to control hearts (*P*=0.0423, **Figure7f-g**). Of note, we also detected an 85% TMPRSS13 increase with I/R injury compared to the control groups (*P=*0.0013, **Figure 7h-i**). These data obtained in the whole heart support the relevance of our findings in NRVM.

### Desmin Oxidation Preserves Cardiac Function with I/R Injury

One obvious PTM induced by oxidative stress is cysteine oxidation, therefore, we sought to test the role of desmin direct oxidation in the models used in this study. As mentioned, there is only one, highly conserved cysteine in the desmin sequence (C332 in *rattus norvegicus*, and C333 in *homo sapiens*). Using maleimide-PEG (MAL-PEG) labeling we measured the oxidation status of this redox-sensitive residue. In short, MAL-PEGylation of free cysteines induces a mobility shift of about ∼ 5kDa which can be detected by classical SDS-PAGE/western blot analysis (**Figure 8a-b**). Using this approach, we were able to measure a ∼22 to ∼24% increase in oxidized desmin at C332 (less available to react with MAL-PEG) in cells treated with 50 µM H_2_O_2_ compared to 10 µM H_2_O_2_ or controls, respectively (both *P*<0.004, **Figure 8b**).

**Figure 8.**
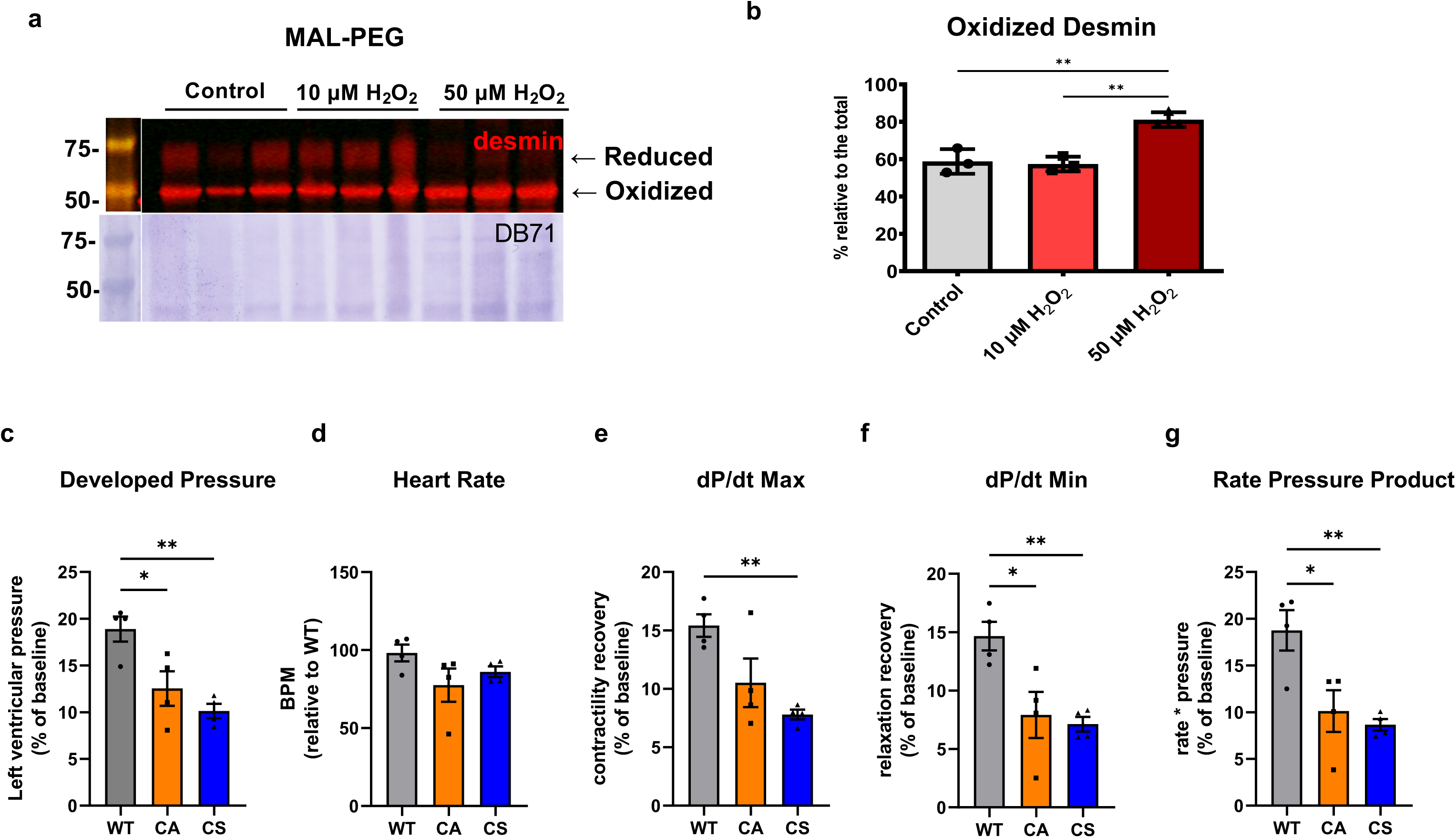
Role of Desmin Oxidation with Oxidative Stress and I/R Injury. **(a)** Representative western blot image for desmin of MAL-PEG-labelled NRVM extracts ± 10 and 50 µM H_2_O_2_ (top); and DB71 for total protein staining (bottom). **(b)** Quantitative measurement of oxidized desmin is relative to total desmin (*n*=3). Quantitative measurements of hemodynamic parameters reflecting functional recovery: **(c)** Developed Pressure; **(d)** Heart Rate; **(e)** systolic function (dP/dt Max); **(f)** diastolic function (dP/dt Min); and **(g)** overall heart function (Rate Pressure Product, RPP) in mouse hearts expressing wild type (WT), C332A (CA) or C332S (CS) desmin. BPM, beat per minute (*n*=4). Mean ± SD is plotted. \*\**P*≤0.01; \**P*≤0.05 by one-way ANOVA followed by Sidak’s multiple comparison test.

To further investigate the functional role of cysteine oxidation with I/R injury, we injected mice with AAV.PHP.s vectors carrying either the wild-type (WT) or redox-dead mimetic (Cys to Ala or CA, and Cys to Ser or CS) desmin sequences. About four weeks after injection hearts were explanted and subjected to I/R as described in **Figure 7a**. We detected a ∼33 to ∼46% drop in left ventricular pressure [Dev P] in CA and CS desmin mutants (*P*=0.0252, *P*=0.004, respectively, **Figure 8c**) compared to WT. While CS mutants also displayed a ∼50% decrease in the recovery of systolic function [dP/dt Max] compared to WT (*P*=0.0079, **Figure 8e**), both CA and CS exhibited a ∼50% drop in the recovery of diastolic function [dP/dt Min] when compared to WT (*P*=0.0181 for CA and *P*=0.0098 for CS, **Figure 8f**). Lastly, there was a ∼50% reduction in overall recovery of heart function [rate pressure product, RPP] for both CA (*P*=0.0214,) and CS (*P*=0.009) compared to WT (**Figure 8g**).

Overall, these data show for the first time that: 1) the simple manipulation of proteoform stoichiometry (as opposed to complete replacement as in knock-in systems) can have measurable, pathophysiological effects; 2) desmin oxidation plays a role in the defense against cardiac oxidative stress. Of note, we could not find a significant difference in desmin cleavage or its aggregation in hearts expressing redox-dead mimetci mutants vs WT desmin. Therefore, while desmin oxidation is protective in the context of I/R injury, the mechanism of this protection seems to differ from a “simple” reduction in loss of gain of desmin function (**Supplementary Figure 2**). The potential signaling roles of desmin’s single cysteine will be addressed in follow-up studies.

## DISCUSSION

There is a compelling need to find strategies that can curb or prevent the acute and long-term adverse effects of reperfusion injury (25). Small molecules that could be administered at the time of percutaneous coronary intervention (PCI) represent ideal candidates and to this end, drugs that elevate *O*-GlcNAc levels with ischemia-reperfusion have shown promise in animal models (8). We reported that the wild-type desmin protein sequence is able to form aggregates, spanning PAOs, fibrillar and large aggregates (4), in various *in vitro* and *in vivo*, small and large animal models of HF as well as human specimens from ischemic and non-ischemic HF patients (5, 6, 20). Notably, our original observation of the amyloidogenic potential of the wild-type desmin sequence was later and independently confirmed by others (10). In fact, Kedia et al., further proposed that desmin aggregates can propagate in a prion-like fashion (10). Because we had reported the accumulation of desmin aggregates in patients suffering from ischemic HF compared to controls (6), we tested here whether desmin aggregates can form as a result of acute oxidative stress that characterizes reperfusion injury. We compared primary cultures of neonatal rat cardiac myocytes and fibroblasts to draw a correlation between desmin aggregation and cell injury induced by oxidative stress. We report here a dose-dependent, cell-type, and IF-protein-specific aggregation of the type III IF desmin (but not of the highly homologous mesenchymal counterpart, vimentin) with acute oxidative stress. Further, desmin aggregation in cardiac myocytes and lack of vimentin aggregation in fibroblasts correlate with cell cytotoxicity, suggesting that desmin aggregation poses a threat for cells that are equipped with lower antioxidant capacity than that of fibroblasts.

New exciting scientific evidence recently highlighted a co-translational role for *O*-GlcNAc in the prevention of protein misfolding and aggregation (7), therefore we tested whether *O*-GlcNAc elevation would efficiently prevent desmin aggregation and cell toxicity in our in vitro model. Indeed, both pre-and post-treatment of cardiac cells subjected to oxidative stress with TMG reduced desmin aggregation and its prodromic PTM (i.e., cleavage), while improving cell survival, in a dose-dependent fashion.

Similar to what we had reported in clinically relevant models and tissue specimens we then confirmed that the accumulation of desmin cleaved forms correlates with damage in the whole heart subjected to I/R injury. One finding from this study whose relevance should not be overlooked is the increase of TMPRSS13 levels in a fashion that mirrors the increase in desmin cleavage. To our knowledge, this is the first report of TMPRSS13 being present at the protein level in cardiac cells and tissue and having a biological function: cleaving desmin. This combined evidence prompts future studies aimed at establishing the cardioprotective potential of TMPRSS13 inhibition with acute and chronic cardiac disease.

Lastly, due to the chemical nature of the pathogenic stimulus used throughout the study, we sought to address the relative contribution of desmin oxidation to its adverse remodeling. As mentioned, type III IF, including desmin, bear only one Cys in their protein sequence that is contained within the 2B coil part of the central, alpha-helical, rod-domain and we postulated that it could work as a sensor for oxidative stress. Using chemical labeling of free cysteines, we confirmed that desmin’s single cysteine is increasingly oxidized in our *in cellulo* model of oxidative stress. Further, using AAV vectors, we demonstrated that co-expression of redox-dead-mimetic desmin (Cys to Ser or Ala), is sufficient to reduce recovery after ischemia-reperfusion injury *ex vivo*. These observations support the hypothesis that desmin oxidation contributes to a protective response against oxidative stress. Intriguingly, our supplementary data show that hearts expressing redox-dead-mimetic desmin mutants have a similar rate of desmin aggregation and cleavage to that of wild-type hearts. This observation suggests that the protective effects provided by desmin’s single cysteine are not directly related to a reduction of desmin gain and/or loss of function. One enticing hypothesis is that desmin’s cysteine could participate in some protective signaling cascade which we will address in follow-up studies.

In all, we provide new evidence for the role of desmin cleavage and its toxic aggregation with cellular oxidative stress and with I/R injury in the working heart. We also firstly suggest that the prevention of desmin aggregation could contribute to the protective role of acute *O*-GlcNAc elevation with ischemia/reperfusion injury, providing a new rationale to design small molecules with more targeted effects and pinpoint a functional role for desmin in the protection from oxidative stress in the heart. While we report the existence and functionality of the new cardiac protease TMPRSS13, which could be potentially targeted to alleviate desmin loss with cardiac injury, we also show that desmin oxidation improves recovery from I/R injury. The relevance and ramification of these last two observations will be addressed in follow-up studies.

## Contributions

ZL, SJ, KKS, PC, and GK performed experiments and contributed to the writing of the experimental portions of the manuscript. KP and HK participated in the experiments. NP provided expertise with the isolated heart models and critical editorial and scientific feedback. GA performed experiments, conceived the study, and wrote the manuscript.

## Acknowledgments

The Authors are grateful to the Zachara’s lab for providing the CTD antibody.

