## Supplementary material for "Ischemia/Reperfusion Injury and Oxidative Stress Impair Cardiac Desmin Proteostasis": Legends to Supplementary Figures

**Supplementary Figure 1. *O*-GlcNAc Elevation by TMG Prior to and Following Oxidative Stress. (a)** Representative western blot image for *O*-GlcNAc (top) and DB71 for total protein staining (bottom) of NRVM pre-treated for 3 hrs and post-treated for 0.5 hr with 2μM and 5μM TMG under 50μM $H_2O_2$. Quantitative measurements of O-GlcNAc level relative to total protein signal for 3 hrs pretreatment **(b)**; and for 0.5 hr posttreatment **(c)** *(n=3)*. Mean ± SD is plotted. **$P$≤0.01; *$P$≤0.05 by two-way ANOVA followed by Sidak's multiple comparison test.

**Supplementary Figure 2. Lack of Changes in Desmin Cleavage and Mono-Phosphorylation with Redox-Dead Mimetic Desmin Mutants Expression. (a)** Representative western blot image for desmin (top) and direct blue 71 (DB71, bottom) for total protein staining. Quantitative measurement of total desmin relative to control **(b)**; desmin cleavage relative to total desmin **(c)**; intact desmin relative to total desmin **(d)**. **(e)** Representative images of phos-tag (top panel) and phos-tag-negative control gels (third panel form the top) for desmin; direct blue 71 (DB71) total protein staining for both blots (second and forth panels from the top). Quantitative measurements for total desmin relative to the control **(f)**; mono-phosphorylation relative to the total **(g)**; tri-phosphorylation relative to the total **(h)** *(n=4)*. Mean ± SD is plotted. **$P$≤0.01; *$P$≤0.05 by one-way ANOVA followed by Sidak's multiple comparison test.
