## Supplementary figures and images for "Ischemia/Reperfusion Injury and Oxidative Stress Impair Cardiac Desmin Proteostasis"

### Supplementary Figures

#### Supplementary Figure 1

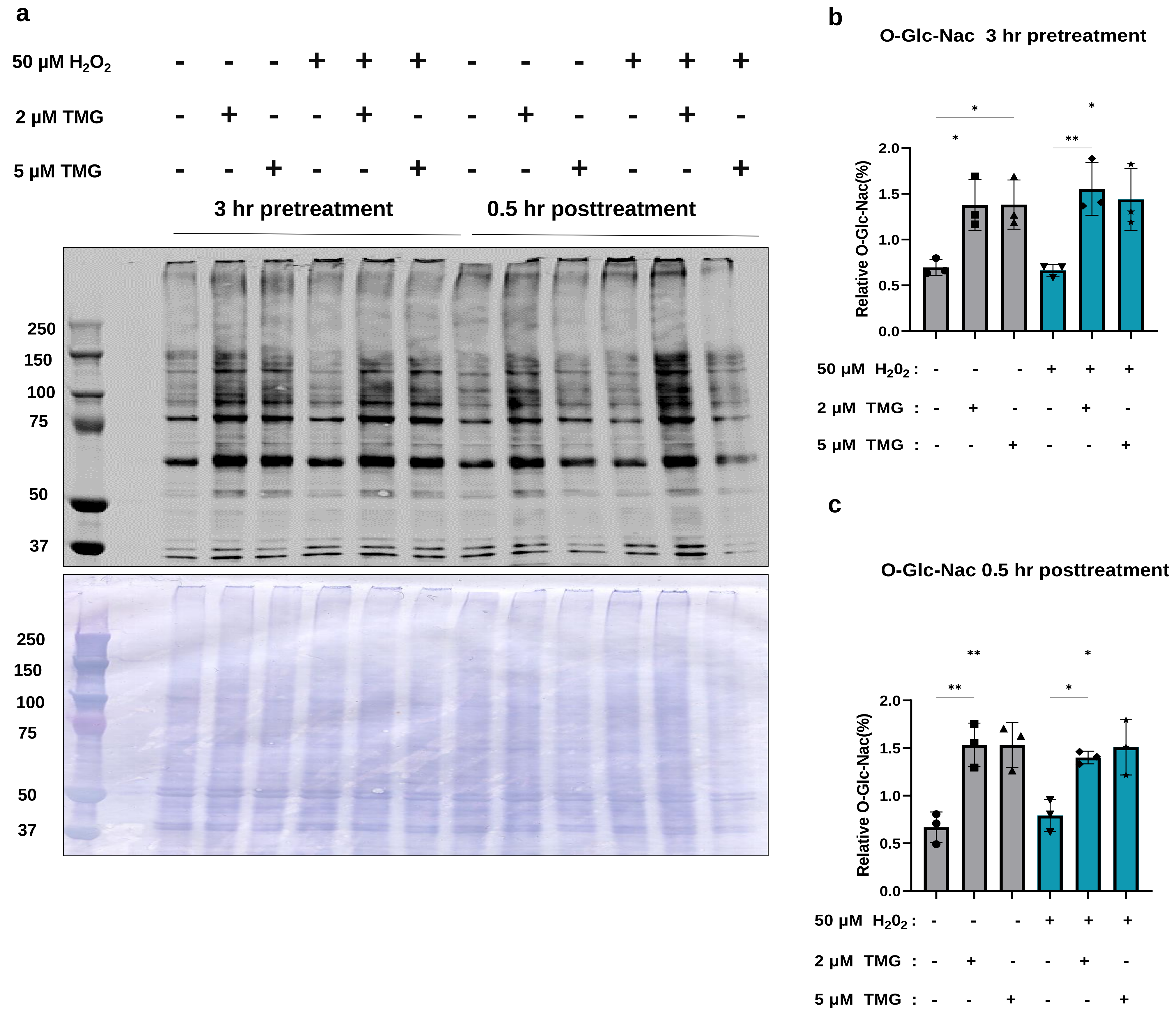

#### Supplementary Figure 2

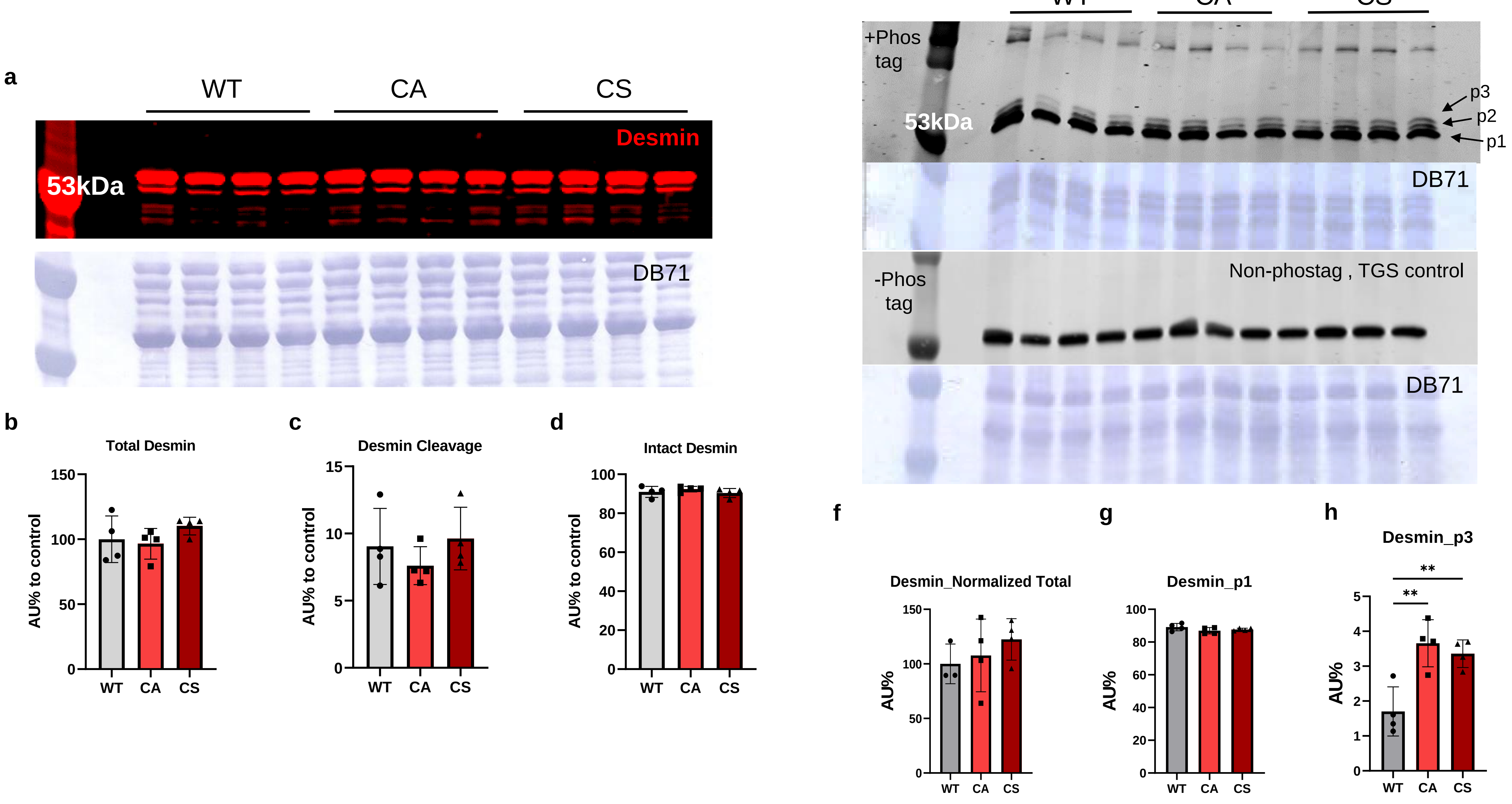
